# Mucosal delivery of a multistage subunit vaccine promotes development of lung-resident memory T cells and affords interleukin-17-dependant protection against pulmonary tuberculosis

**DOI:** 10.1101/2020.02.25.964312

**Authors:** Claudio Counoupas, Kia Ferrell, Anneliese Ashhurst, Nayan Bhattacharyya, Gayathri Nagalingam, Carl G. Feng, Nikolai Petrovsky, Warwick J. Britton, James A. Triccas

## Abstract

The development of effective vaccines against bacterial lung infections requires the induction of protective, pathogen-specific immune responses without deleterious inflammation within the pulmonary environment. Here, we made use of a polysaccharide-adjuvanted vaccine approach to elicit resident pulmonary T cells to protect against aerosol *Mycobacterium tuberculosis* infection. Intratracheal administration of the multistage fusion protein CysVac2 and the delta-inulin adjuvant Advax™ (formulated with a TLR9 agonist) provided superior protection against aerosol *M. tuberculosis* infection in mice, compared to parenteral delivery. Surprisingly, removal of the TLR9 agonist did not impact vaccine protection despite a reduction in cytokine-secreting T cell subsets, particularly CD4^+^ IFN-γ^+^IL-2^+^TNF^+^ multifunctional T cells. CysVac2/Advax-mediated protection was associated with the induction of lung-resident, antigen-specific memory CD4^+^ T cells that expressed IL-17 and RORγt, the master transcriptional regulator of Th17 differentiation. IL-17 was identified as a key mediator of vaccine efficacy, with blocking of IL-17 during *M. tuberculosis* challenge reducing phagocyte influx, suppressing priming of pathogen-specific CD4^+^ T cells in local lymph nodes and ablating vaccine-induced protection. These findings suggest that tuberculosis vaccines such as CysVac2/Advax that are capable of eliciting Th17 lung-resident memory T cells are promising candidates for progression to human trials.

**Importance:** *Mycobacterium tuberculosis*, the causative agent of tuberculosis (TB), kills more individuals each year than any other single pathogen. The only approved vaccine, BCG, administered intradermally, is unreliable in protecting against pulmonary TB, therefore a more effective vaccine is critical for global control of the disease. Vaccination in the lung would be a rational way of inducing a local memory immune response to TB, however vaccine platforms would need to deliver antigens to delicate mucosal surfaces without inducing deleterious inflammatory responses. We developed a safe mucosal vaccine which induced protection against TB lung infection in mice by inducing high levels of lung-resident T cells expressing the cytokine IL-17. Removal of IL-17 limited the influx of phagocytic cells to the lung and completely ablated protection afforded by the vaccine. This study provides new insights into mechanisms of protection against *M. tuberculosis* and provides a promising candidate to protect against TB in humans.

## Introduction

Tuberculosis (TB) remains a major cause of morbidity and mortality worldwide, with 10 million new cases and 1.7 million deaths per year[1]. *Mycobacterium bovis* bacillus Calmette-Guérin (BCG) is currently the only licensed vaccine against TB, however its efficacy varies greatly, especially against the adult pulmonary form of the disease[2]. The 2015 WHO End TB Strategy identified the development of a more effective and easily administered vaccine for controlling TB and halting the global epidemic[3]. In recent decades, extensive research has resulted in many new TB vaccine candidate, 14 of which are currently in human vaccine trials and are reviewed in detail elsewhere[4]. Recently, a Phase IIb clinical trial of the fusion protein vaccine M72/AS01_E_ showed protective efficacy of 50% in *M. tuberculosis*-infected adults after 3 years [5, 6]. Although promising, vaccines with higher efficacy are considered necessary to reduce TB incidence to the targets outlined in the End TB Strategy objectives[3].

One of the major limitations of current vaccination strategies is that the administration route may not be optimal for the induction of immunity at the site of pathogen entry, i.e. the lung. Pulmonary vaccine delivery has been hindered by the fact that most adjuvants are either unable to induce sufficient mucosal immunity or are too toxic to be administered to the lung[7]. However, recent evidence supports the idea that mucosal vaccination may provide superior protection against respiratory *M. tuberculosis* infection over parenteral vaccination. For example, lung-resident CD4^+^ memory T cells (T_RM_) induced after pulmonary vaccination with a recombinant influenza virus expressing *M. tuberculosis* antigens provided protection in the lung in the absence of circulating memory cells[8]. T_RM_ have also been proposed as the possible mechanism of protection in macaques that demonstrate sterilizing immunity after intravenous vaccination with BCG and subsequent *M. tuberculosis* infection [9]. When administered through the mucosal route, BCG induced increased protection compared to the intradermal immunization, which was linked to lung T_RM_ and Th17 polarization of the CD4^+^ T cells[10]. Th17 responses have been associated with the influx of neutrophils with bactericidal activity[11] and increased CD4^+^ T cell recruitment to the lung after *M. tuberculosis* infection[12]. Vaccines inducing high levels of pulmonary IL-17 have demonstrated efficacy against *M. tuberculosis* in different animal models[13, 14] although balancing the protective and pathogenic roles of IL-17 in the lung is a critical consideration[15].

In this study we sought to determine if the candidate TB vaccine, CysVac2/Advax[16], is effective as a mucosal vaccine to protect against *M. tuberculosis.* CysVac2 is a fusion protein of two *M. tuberculosis* antigens; the immunodominant Ag85B and CysD, a component of the sulfur assimilation pathway that is overexpressed in chronic stages of infection [17]. Advax is a particulate polysaccharide adjuvant with a low inflammatory profile that has proven to be safe and a strong inducer of vaccine immunogenicity in humans, thus making it an ideal candidate for mucosal administration[18][19]. Notably, it was recently shown to provide safe and effective enhancement of influenza vaccine immunity when administered via the intrapulmonary route in different animal models[20, 21].

We report here that intrapulmonary administration of CysVac2/Advax induced greater protection in mice than parenterally administered vaccine, with the vaccine promoting the accumulation of antigen-specific, IL-17-secreting CD4^+^ T_RM_ in the lungs. Furthermore, IL-17 was essential for the protective efficacy afforded by intrapulmonary CysVac2/Advax vaccine, thus defining a crucial role for this cytokine in vaccine-mediated control of TB.

## Results

### Pulmonary administration of CysVac2/Advax^CpG^ provides superior protection against *M. tuberculosis* challenge than parenteral vaccination

Previous studies of intramuscular (i.m.) vaccination of mice with CysVac2/Advax^CpG^ demonstrated substantially enhanced systemic CD4^+^ T cell responses composed of multifunctional Th1 polarized cells, which correlated with protection against aerosol *M. tuberculosis* infection[16]. In this study, we evaluated if delivery to the lung by intratracheal (i.t.) instillation of this vaccine candidate could improve the level of protection induced by this vaccine. Mice were vaccinated by either the i.t. or i.m. routes with CysVac2/Advax^CpG^ 3 times, 2 weeks apart (Fig 1a). When the vaccine-specific T cell response was examined in the blood prior to *M. tuberculosis* challenge, a higher level of circulating polyfunctional CD4^+^ T cells expressing IFN-γ were present after i.m. vaccination (Fig 1b, S1a Fig), with the most prominent phenotype identified as multi-cytokine secreting CD44^+^ CD4^+^ T cells (Fig 1c). After i.t vaccination with CysVac2/Advax^CpG^, PBMC-derived CD4^+^ T cell expressing either IL-2, TNF or IL-17 were more prominent when compared to the i.m route (Fig 1b). Both vaccination regimens induced similar proportions of T-bet expression in circulating CD4^+^ T cells, however i.t. vaccination induced a higher proportion of cells expressing RORγT (Fig 1d).

**Figure 1.**
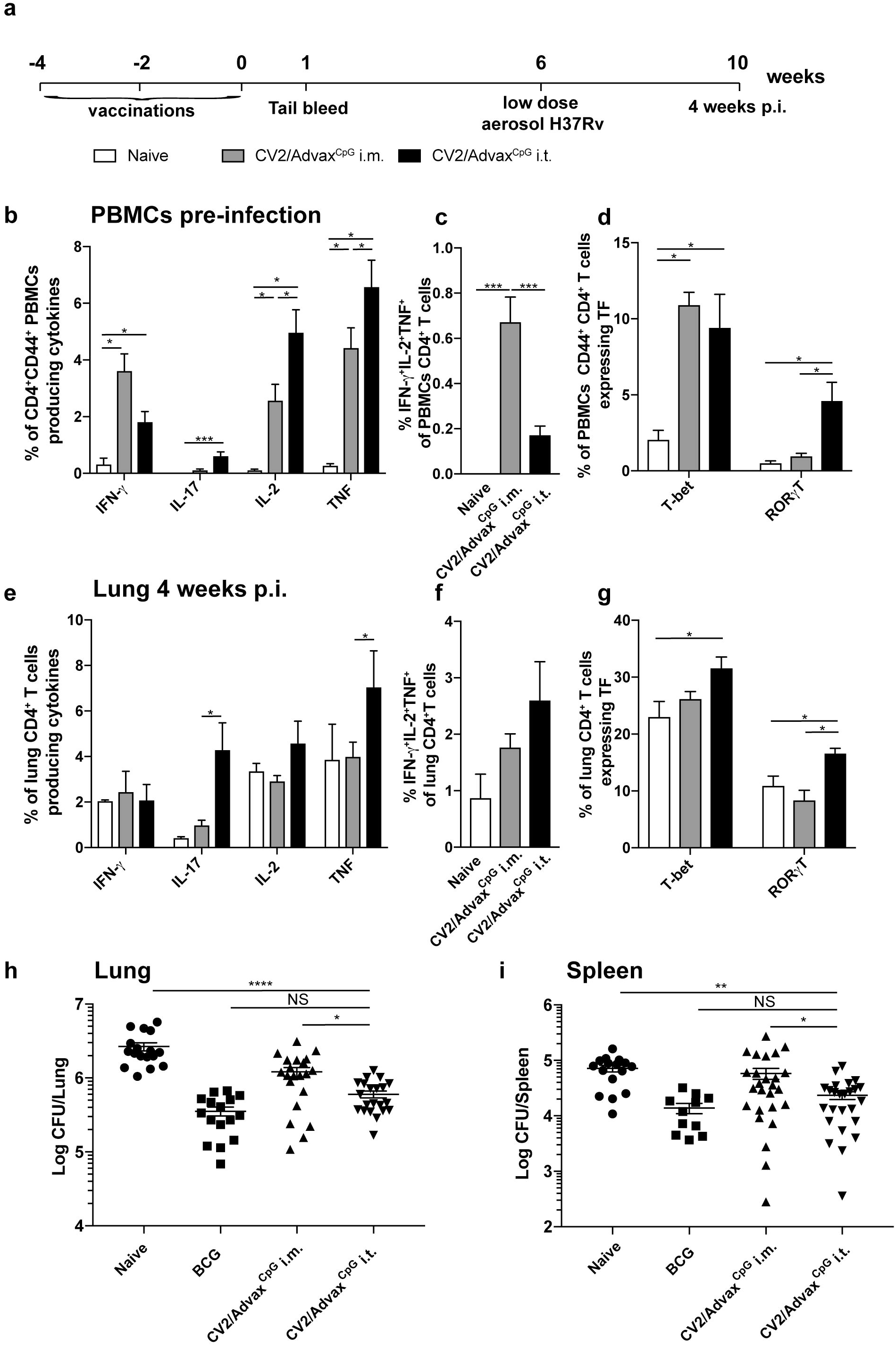
Pulmonary vaccination with CysVac2/Advax^CpG^ demonstrates improved protection against *M. tuberculosis* infection compared to parenteral administration. C57BL/6 mice (n=5-6) were vaccinated by either the i.m. or i.t. route with CysVac2(CV2)/Advax^CpG^ (3 times, 2 weeks apart). One week after last vaccination mice were bled for vaccine immunogenicity assessment. Six weeks after last immunization mice were challenged with H37Rv by aerosol (~100 CFU) and four weeks later culled to enumerate bacterial burden and T cell phenotype in the lung (a). PBMCs from tail blood of vaccinated mice (b, c, d) or cells from lung of infected mice (e, f, g) were restimulated *ex vivo* with CysVac2, and the production cytokines (IFN-γ, IL-2, IL-17, TNF), or transcription factors (TF; T-bet, RORγT) by CD4^+^ T cells was determined by flow cytometry. Data are represented as the percentage of cytokine-producing or transcription factor-positive CD4^+^ T cells ± SEM. Bacterial load was assessed in the lungs (h) and in the spleen (i) and presented as Log_10_ of the mean CFU ± SEM. Data are pooled from 3 independent experiments. Significance of differences between the groups was determined by ANOVA (*p<0.05; **p<0.01; ***p<0.001).

The pattern of CD4^+^ T cells immune responses pre-*M. tuberculosis* challenge was compared to that after *M. tuberculosis* infection. In i.m. vaccinated mice, the greatest frequency of CD4^+^ T cells observed were those secreting IFN-γ or TNF (Fig 1e) and this was dominated by cells with a polyfunctional Th1 responses (IFN-γ^+^IL-2^+^TNF^+^, Fig 1f). I.t. vaccination with CysVac2/Advax^CpG^ resulted in a high frequency of CD4^+^ T cells secreting either IL-17 or TNF (Fig 1e) or both cytokines (S1b Fig); these T cell subsets were not observed after i.m. vaccination (Fig 1e, S1 Fig). Further, the frequency of CD4^+^ T cells expressing RORγT was significantly enhanced after i.t. vaccination compared to unvaccinated or i.m. vaccinated mice (Fig 1g). Therefore i.t. vaccination of mice with CysVac2/Advax^CpG^ results in a Th17-polarized T cell response post-*M. tuberculosis* exposure, which was not observed after i.m. immunization.

Considering the differential pattern of immune responses induced by varying the route of administration of CysVac2/Advax^CpG^, we next determined if this had an impact on protective efficacy. I.t. vaccinated mice challenged with low dose aerosol *M. tuberculosis*, demonstrated significantly enhanced lung protection when compared to i.m.-vaccinated or unvaccinated mice (Fig 1h). A similar result was observed in the spleen, suggesting that i.t. vaccination might improve protection against disseminated infection (Fig 1i). Taken together, these results demonstrate that pulmonary vaccination with CysVac2/Advax^CpG^ induces superior protection compared to i.m. vaccination and this is associated with an enhanced generation of Th17 cells in the circulation and in the lung.

### CpG is dispensable for protection generated by pulmonary vaccination with CysVac2/Advax

CpG oligonucleotides are TLR9 agonists that help drive Th1 immune responses[22]. While the CpG component was shown to be important to the protection obtained after CysVac2 i.m. immunization, we were interested whether a simplified formulation of Advax without the CpG component would still generate protective pulmonary immunity. CysVac2/Advax or CysVac2/Advax^CpG^ were delivered by the i.t. route, and the mice challenged with aerosol *M. tuberculosis*. Both vaccines resulted in the generation of IL-17-producing CD4^+^ T cells following antigen restimulation *ex vivo* (Fig 2A), however all inflammatory cytokines (IFN-γ, IL-17, TNF) were reduced in Advax-immunized as compared to Advax^CpG^-vaccinated mice (Fig 2a). Strikingly, the removal of the CpG component resulted in the loss of multifunctional CD4^+^ T cells with a triple cytokine-secreting profile (IFN-γ^+^IL-2^+^TNF^+^) (Fig 2b). However, both vaccinated groups displayed equivalent expression of the transcription factors T-bet or RORγT (Fig 2c) and similar protection against *M. tuberculosis* in the lungs (Fig 2c) and spleen (Fig 2d). This indicates that CysVac2/Advax is sufficient for protection and this protection does not correlate with the presence of multifunctional T cells secreting high levels of inflammatory cytokines.

**Figure 2.**
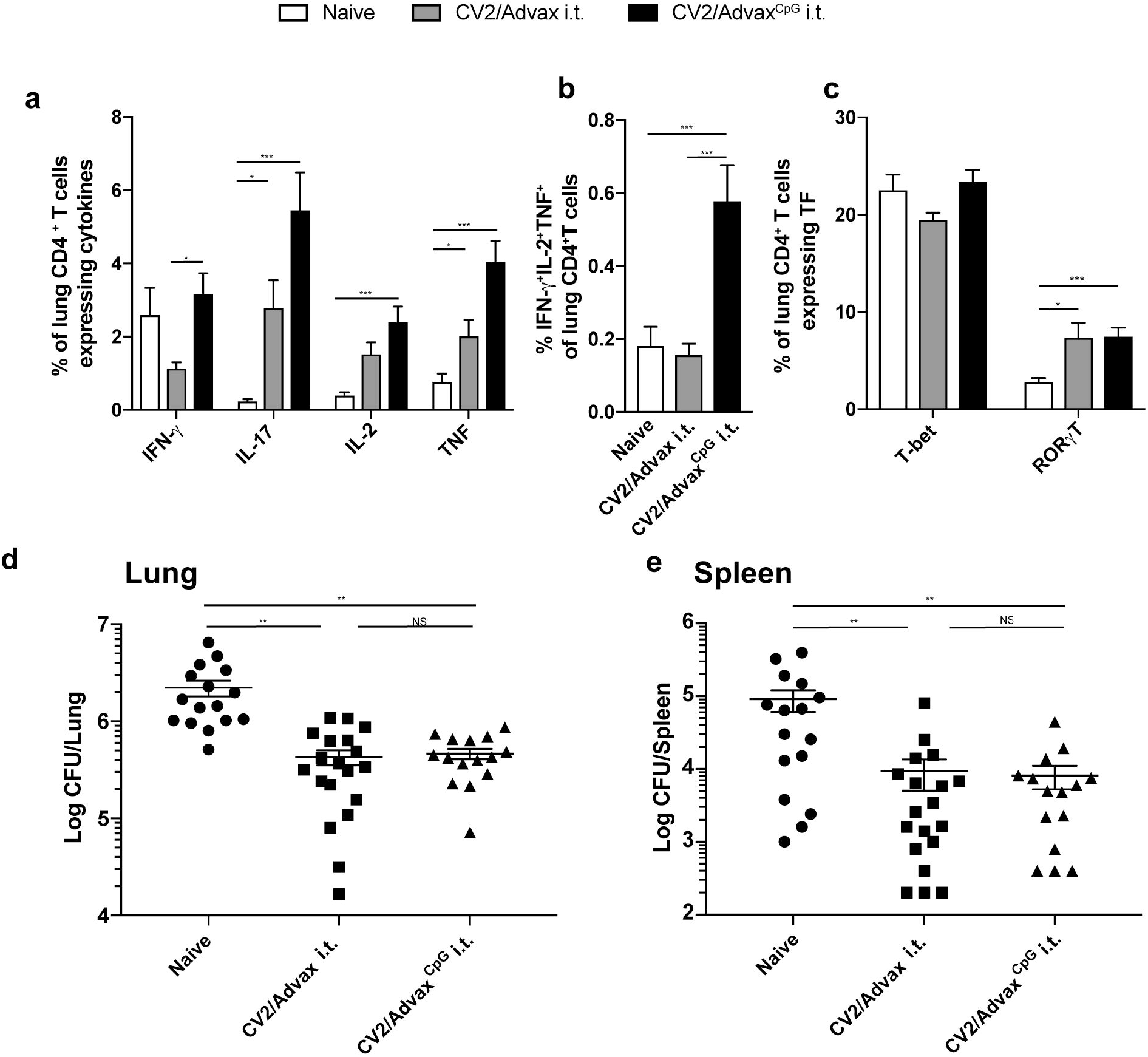
CpG is dispensable for protective immunity induced by CysVac2/Advax. C57BL/6 mice (n=5-6) were vaccinated by the i.t. route with CysVac2/Advax^CpG^ or CysVac2/Advax (3 times, 2 weeks apart). Six weeks after last immunization mice were challenged with *M. tuberculosis* H37Rv by aerosol (~100 CFU) and four weeks later culled to enumerate bacterial burden and T cell phenotype in the lung. Cells from lung of infected mice were restimulated *ex vivo* with CysVac2 and the production of cytokines (IFN-γ, IL-2, IL-17, TNF; panel a and b) or transcription factors (T-bet, RORγT; panel c) by CD4^+^ T cells was determined by flow cytometry. Data are represented as the percentage of cytokine-producing CD4^+^ T cells ± SEM. Bacterial load was assessed in the lungs (d) and in the spleen (e) and presented as Log_10_ of the mean CFU ± SEM. Data are pooled from 2 independent experiments. Significance of differences between the groups was determined by ANOVA (*p<0.05; **p<0.01; ***p<0.001).

### Intrapulmonary CysVac2/Advax generates lung-resident, antigen-specific CD4^+^ T cells

To more precisely define the vaccine-specific responses after immunization with CysVac2/Advax, Ag85B:I-A^b^ tetramer staining was employed to identify CD4^+^ T cells specific for the p25 epitope of Ag85B, an antigenic component of CysVac2[17]. Ag85B tetramer-positive (Ag85Btet^+^) cells in the lungs were only detected after i.t. delivery of CysVac2/Advax and not after CysVac2 antigen alone, confirming the critical role of Advax in inducing antigen-specific T cell expansion (Fig 3a). Ag85Btet^+^ cells were present in significantly greater numbers in CysVac2/Advax vaccinated samples at all timepoints examined, although numbers contracted by 8 weeks post-vaccination (Fig 3b). CysVac2/Advax-vaccinated mice showed enrichment of lung parenchymal-residing (intravascular negative IV^−^) CD4^+^ T cells, which expressed CD69 and CD44 with low level of expression of the lymphoid homing receptor, L-selectin (CD62L) (Fig 3d). This population of CD4^+^CD44^hi^CD62L^low^CD69^+^ IV^−^ were defined as T_RM_-like cells[23] and were significantly greater at all times points post-vaccination in CysVac2/Advax-vaccinated mice, comprising approximately 25% of total CD4^+^ T cells in the lung following vaccination (Fig 3c). The majority of Ag85B-tet^+^ cells detected at 8 weeks post vaccination showed a T_RM_-like phenotype and were present within the parenchyma of the lung (Fig 3e). Along with T_RM_ markers, this subset expressed high levels of the integrin CD11a and the cell surface receptor PD-1, with low levels of expression of CD103 and KLRG-1 (Fig 3e). Taken together, these data indicate that CysVac2/Advax vaccination induces a population of antigen-specific CD4^+^ T cells in the lung with a T_RM_ phenotype, and these are detectable through 8 weeks post vaccination.

**Figure 3.**
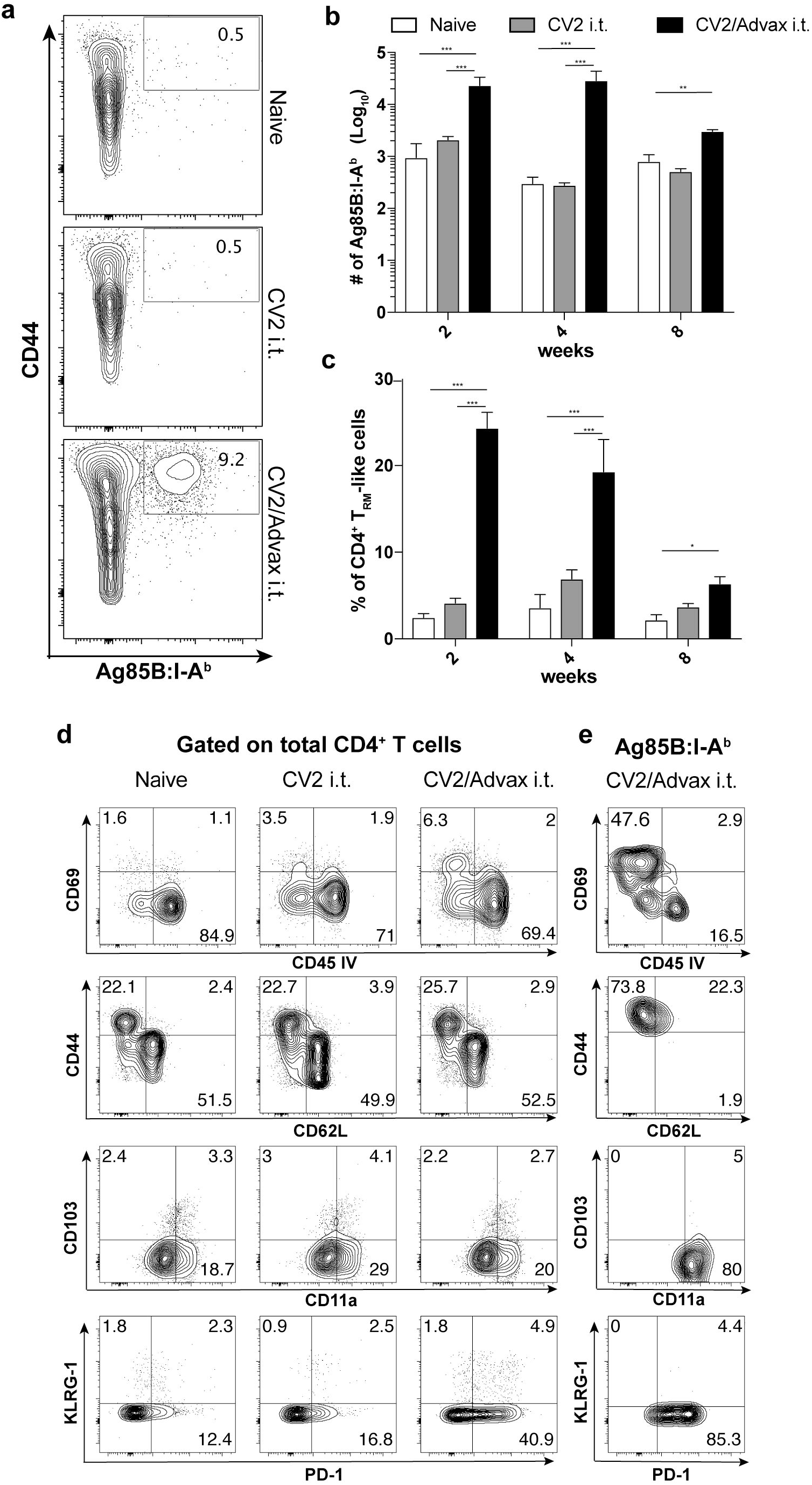
Pulmonary vaccination with CysVac2/Advax induces Ag-specific persistent local resident CD4^+^ T cells. C57BL/6 mice (n=3-4) were vaccinated by the i.t. route with CysVac2/Advax or CysVac2 (3 times, 2 weeks apart). At weeks 2, 4, or 8 after final immunization, lung cells were processed for Ag85B:I-A^b^ tetramer staining (a, Representative dot plot from 8 weeks post-immunization). Number of Ag85B:I-A^b^ tetramer positive cells in the lung over time are shown in (b). Also shown are the percentage of total CD4^+^ T cells expressing phenotypic makers markers associated with TRMs (CD45 IV^−^, CD11a^+^, CD69^+^, CD44^+^, PD1^+^ KLRG^−^, c) and representative dot plots of TRMs markers on total lung CD4^+^ T cells (d) or in Ag85B:I-A^b+^ cells (e) at 8 weeks after last vaccination. Data are representative of 2 independent experiments. Significance of differences between the groups was determined by ANOVA (*p<0.05; **p<0.01; ***p<0.001).

We further characterized the ability of the CD4^+^ T cells generated in the lung to produce cytokines in response to re-stimulation with vaccine antigen at time points after vaccination (pre-challenge) as well as at 4 weeks post-challenge with *M. tuberculosis*. Intratracheal CysVac2/Advax vaccination induced a distinct cytokine profile that was relatively consistent across all time points prior to *M. tuberculosis* infection. While a marginal increase in IFN-γ production was observed at 4 weeks post vaccination, this was not apparent at any other time point (Fig 4a). Indeed, following infection with *M. tuberculosis*, a lower percentage of IFN-γ-producing CD4^+^ T cells was present in CysVac2/Advax vaccinated lung samples compared with samples from unvaccinated animals. By contrast, percentages of of IL-2, IL-17 or TNF cytokine-producing cells were markedly increased after i.t. vaccination with CysVac2/Advax compared to unvaccinated or CysVac2 vaccinated mice up to 8 weeks post-vaccination (Fig 4b, 4c, 4d), however only IL-17 remained elevated post-challenge (Fig 4b). Further analysis revealed that the major source of IL-17 originated from CD45 IV^−^CD4^+^ T cells that expressed RORγT (Fig 4e). Furthermore, the Ag85Btet^+^ population was highly enriched in RORγT^+^ CD45 IV^−^IL-17^+^CD4^+^ T cells (Fig 4f). Overall, these data suggest that pulmonary vaccination with CysVac2/Advax promotes increased single and multifunctional cytokine producing T cell populations both before and after aerosol challenge with *M. tuberculosis*, which is characterized by the development of a tissue-resident Th17-type response.

**Figure 4.**
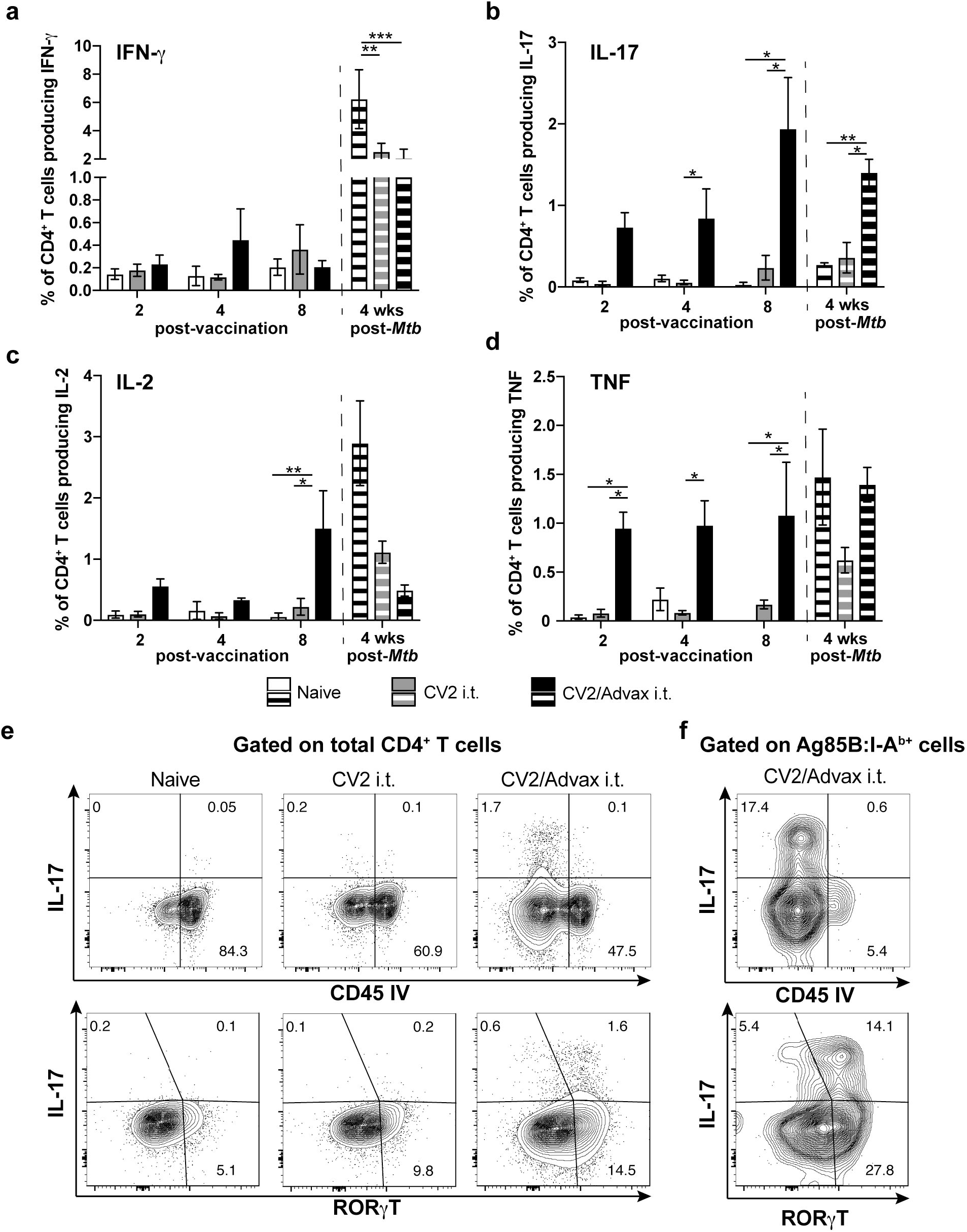
Persistent CysVac2-specific IL-17 production by lung-resident CD4^+^ T cells after pulmonary vaccination with CysVac2/Advax. C57BL/6 mice (n=4) were vaccinated by the i.t. route with CysVac2/Advax or CysVac2 protein alone (3 times, 2 weeks apart). Eight weeks after the last immunization mice were challenged with *M. tuberculosis* H37Rv by aerosol (~100 CFU). At 2, 4, or 8 weeks after last immunization (solid bars), and at 4 weeks after infection (striped bars), lung cells were restimulated *ex vivo* with CysVac2 and the production of IFN-γ (a), IL-17 (b), IL-2 (c) or TNF (d) by CD4^+^ T cells determined by flow cytometry. Representative dot plots of co-expression of CD45 IV or RORγT with IL-17 by total CD4^+^ T cells (e) or Ag85B:I-A^b^ tetramer positive cells (f) at 8 weeks after last vaccination. Data are represented as the percentage of cytokine-producing CD4^+^ T cells ± SEM and is representative of 2 independent experiments. Significance of differences between the groups was determined by ANOVA (*p<0.05; **p<0.01; ***p<0.001).

### IL-17-mediated protection correlates with early recruitment of phagocytic cells and enhanced priming of pathogen-specific CD4^+^ T cells

Given the marked Th17 polarization of the CD4^+^ T cell response to pulmonary immunization with CysVac2/Advax, we next determined the impact of neutralizing IL-17 at the time of *M. tuberculosis* challenge (Fig 5a). Treatment with anti-IL-17 mAb did not affect the capacity of CD4^+^ T cells to respond to infection, as the frequency of cytokine-producing CD4^+^ T cells was not altered between mice treated with anti-IL-17 or isotype control mAb (Fig 5b). However, anti-IL-17 treatment had a detrimental effect on control of bacterial infection in the lung of CysVac2/Advax immunised mice; bacteria numbers in anti-IL-17 treated mice were similar to unvaccinated mice, while immunised mice treated with isotype control mAb remained protected against infection (Fig 5c).

**Figure 5.**
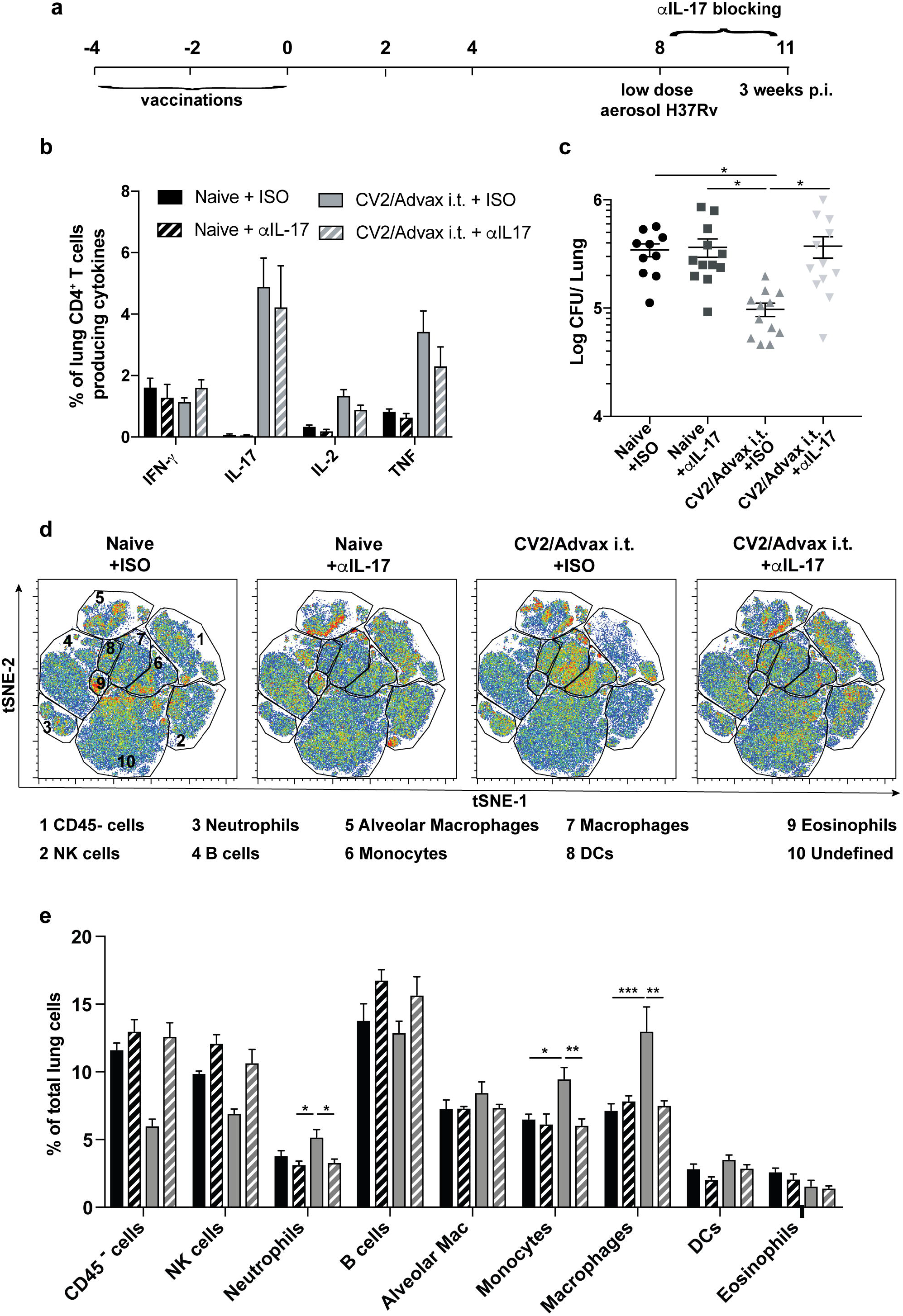
Protection afforded by pulmonary CysVac2/Advax against aerosol *M. tuberculosis* is dependent on IL-17 and correlates with lung phagocytic cells recruitment. C57BL/6 mice (n=6) were vaccinated by the i.t. route with CysVac2/Advax (3 times, 2 weeks apart) and at 8 weeks after last immunization mice were challenged with *M. tuberculosis* H37Rv by aerosol (~100 CFU). One day before aerosol, mice were treated i.p. with anti-IL1-7 or an isotype control mAb (twice weekly for 3 weeks) (a). Cells from the lungs of infected mice were restimulated *ex vivo* with CysVac2 and cytokines secretion (IFN-γ, IL-2, IL-17, TNF) determined (b). Bacterial load was assessed in the lungs and is presented as Log_10_ of the mean CFU ± SEM (c). Representative tSNE dimension 1 and 2 plots of the total live cells in the lung (d). Bar graphs show the percentage of identified lung cells subsets (e). Data are pooled of 2 independent experiments. Significance of differences between the groups was determined by ANOVA (*p<0.05; **p<0.01; ***p<0.001).

We also studied the impact of IL-17 blocking on lung cell subsets after *M. tuberculosis* infection using flow cytometry phenotyping combined with an unsupervised visual implementation of t-distributed stochastic neighbour embedding (tSNE) analysis. The generated tSNE plot was calculated with 12 parameters and 10 clusters obtained using unsupervised analysis were subsequently assigned to a specific cell population (Fig 5d), according to expression level of each marker and previously described phenotypes (S3Fig). This analysis revealed that the percentage of neutrophils in the lung were elevated in vaccinated mice treated with control mAb, however they returned to the level of unvaccinted animals after treatment with the anti-IL-17 mAb (Fig 5e). A similar pattern was observed for monocytes and monocyte-derived-macrophages, however no differences were observed for other phagocytic populations in the lung, such as alveolar macrophages (Fig 5e).

We next investigated the effect of blocking IL-17 on the priming and proliferation of CD4^+^ T cells. To do this, we examined CD4^+^ T cells primed by the vaccination (Ag85B-tet^+^) and compared to T cells responding specifically to *M. tuberculosis* (ESAT6-tet^+^). In the lung we observed a greater proportion of Ag85B-tet^+^ cells in vaccinated mice compared to unvaccinated mice, however anti-IL-17 treatment did not significantly alter the numbers of proliferating CD4^+^ T cells (S4a, b, and c Fig) or the numbers of Ag85B-tet^+^ or ESAT6-tet^+^ cells (S4d, e and f Fig). However, in the mLN of vaccinated mice a distinct vaccine-induced population of proliferating CD4^+^ T cells expressing RORγT was distinguishable (Fig 6a). IL-17 blocking resulted in reduced proliferation of total CD4^+^ T cells in the mLN of vaccinated mice (Fig 6b) including the vaccine-primed RORγT^+^ cells (Fig 6c). When the frequency of the vaccine-primed CD4^+^ T cells (Ag85B-tet^+^) were compared to those primed only by *M. tuberculosis* infection (ESAT6-tet^+^) (Fig 6d), Ag85B-tet^+^ cells were significantly higher in vaccinated mice compared to the unvaccinated group, however anti-IL-17 treatment did not affect the numbers of Ag85B-tet^+^ cells (Fig 6e). By contrast, the numbers of ESAT6-tet^+^ positive CD4^+^ T cells were reduced after IL-17 blocking in the mLNs (Fig 6f). Overall, neutralizing of IL-17 reduced phagocytic cells in the lung of immunized animals and the numbers of pathogen-specific CD4^+^ T cells in the draining LNs, which correlated with a loss of vaccine-mediated protection.

**Figure 6.**
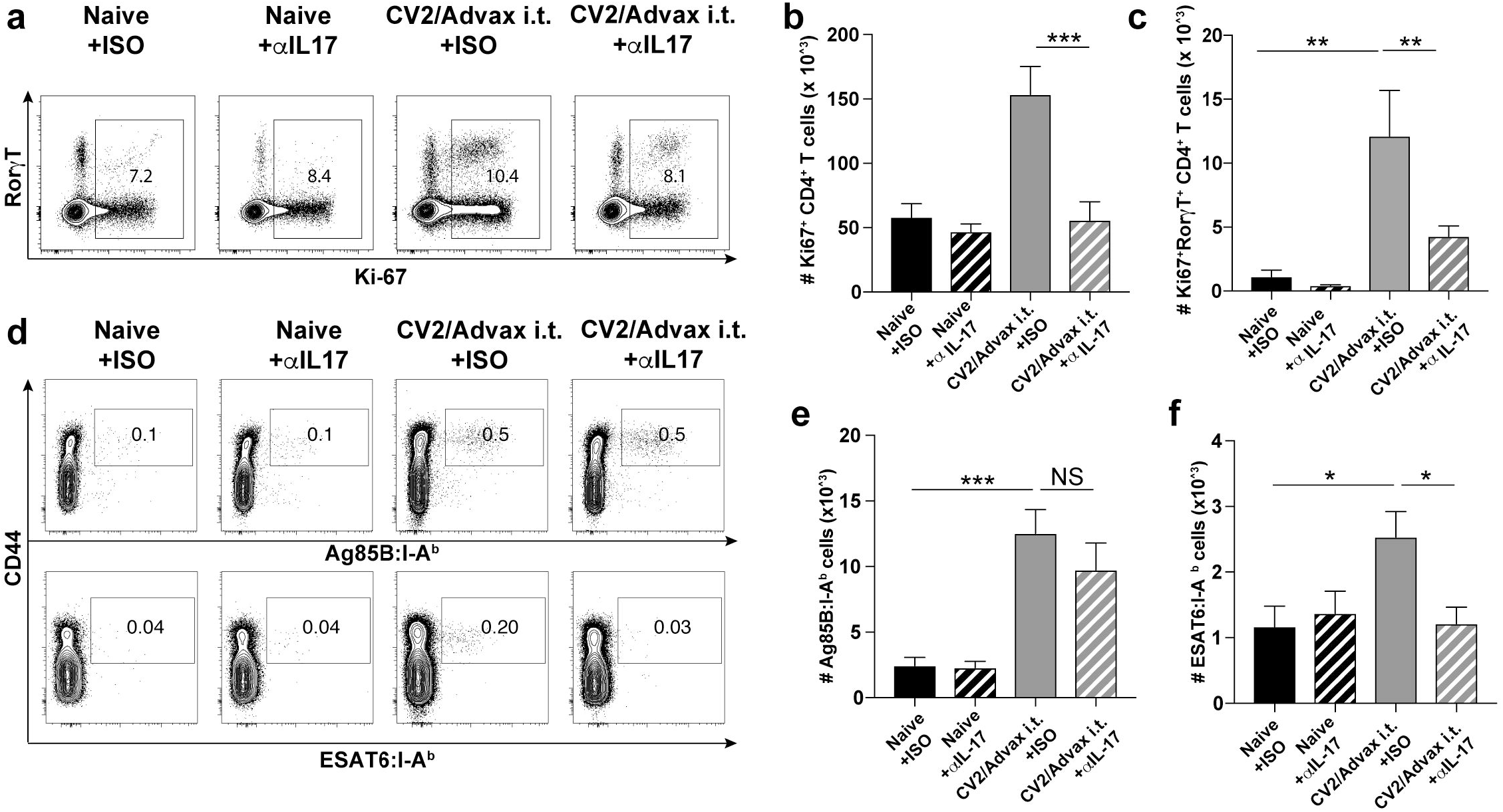
Blocking IL-17 during *M. tuberculosis* infection impairs the proliferation of pathogen-specific CD4^+^ T cells in the mediastinal lymph nodes. C57BL/6 mice (n=6) were vaccinated with CysVac2/Advax and treated i.p. with anti-IL-17 mAb, as described in Figure 5. Representative dot plot of the expression of Ki67 and RORγT on CD4^+^ T cells (a). Bar graphs show numbers of total (b) and RORγT^+^ (c) proliferating CD4^+^ T cells enumerated in the mLN. Representative dot plots show CD44 and either Ag85B:I-A^b^ or ESAT6:I-A^b^ staining on CD4^+^ T cells in the mLN (d), with total number ± SEM of Ag85B:I-A^b+^ (e) and ESAT6:I-A^b+^ (f) CD4^+^ T cells in the mLN. Data are pooled of 2 independent experiments. Significance of differences between the groups was determined by ANOVA (*p<0.05; **p< 0.01).

## Discussion

The respiratory tract is the preferred port of entry of *M. tuberculosis*, with the complexity of the immunological environment in the lung potentially contributing to suboptimal pathogen responses. This may be particularly detrimental in the case of *M. tuberculosis* infection, where priming and recruitment of effector T lymphocytes to the lungs is delayed, allowing unchecked growth of the organism[24, 25]. For this reason, mucosal vaccination has been of interest in the field of TB vaccines, with pulmonary delivery of BCG[14, 26], live recombinant viruses[8, 27, 28], or protein/adjuvants[29, 30] resulting in protective immune responses. When administered to the respiratory mucosa, highly inflammatory adjuvanted vaccines may induce protective immunity, but this is often accompanied by excessive inflammation, mucus accumulation and eosinophilia[31]. Thus, there is a need for adjuvants that can induce protective lung immunity in the absence of deleterious inflammation and pathology. We found that i.t. delivery of CysVac2/Advax^CpG^ was significantly more protective than i.m. vaccination against challenge with *M. tuberculosis* (Fig 1). Strikingly i.t. vaccination, despite being more protective, did not induce appreciable numbers of multifunctional CD4^+^ T cells (IFN-γ^+^IL-2^+^TNF^+^; Fig 1) and even more surprisingly removal of CpG from the formulation did not reduce vaccine protection, despite a loss of multifunctional CD4^+^ T cell generation (Fig 2). Induction of multifunctional T cell responses has been used as a key criteria for vaccine progression to human trials, however recent evidence in both mice and humans indicates that the generation of IFN-γ secreting T cells does not necessarily correlate with protection[32]. Indeed, in our data IFN-γ was the only cytokine analyzed pre- and post-challenge that did not correlate with the protective effect of the CysVac2/Advax vaccine (Fig 4), an important finding for the selection of vaccines for progression to human trials.

We observed that pulmonary vaccination with CysVac2 vaccination induced a lung resident CD4^+^ population that expressed markers of T_RM_-like cells (Fig 3). We have previously shown that a recombinant influenza vaccine conferred protection against *M. tuberculosis* in the absence of circulating memory T cells, suggesting an important role for *M. tuberculosis*-specific T_RM_[8]. In the current study, detailed phenotypic analysis of CysVac2/Advax-induced, antigen-specific CD4^+^ T_RM_-like cells showed that in addition to well-characterized markers of tissue-resident memory cells (CD69^+^ CD44^hi^ CD62L^low^ CD45 IV^−^), this population displayed high levels of CD11a with minimal expression of CD103 (Fig 3). This low level of CD103 expression contrasts with the increased expression of CD103 on CD8 T_RM_, where this integrin is thought to be essential in the retention of these cells within tissue[33]. Antigen-specific T_RM_ cells induced following CysVac2/Advax vaccination also displayed a dominant PD-1^+^ KLRG-1^−^ phenotype (Fig 3). During *M. tuberculosis* infection, PD-1 expression indicates an earlier stage of CD4^+^ T cell differentiation that is associated with a higher proliferative capacity, while KLRG-1^+^ cells are more terminally differentiated and produce greater levels of cytokines[34]. IL-2-secreting KLRG-1^−^ T cells induced by subunit booster vaccination are associated with protection against chronic *M. tuberculosis*, owing to maintenance of their proliferative capacity[35], and circulating KLRG-1^−^ CD4^+^ T cells induced by subcutaneous H56/CAF01 vaccination display the ability to home to the lungs[36]. Our findings confirmed an association between vaccine-induced, PD-1^+^KLRG-1^−^ CD4^+^ T cells that reside within the lung parenchyma and protection against *M. tuberculosis*. It is possible these cells possess greater effector functions and proliferative capacity than their KLRG-1^+^ counterparts and are therefore capable of enhanced protection against *M. tuberculosis* infection.

Lung-resident CD4^+^ T_RM_ were found to be an important source of IL-17 induced after Advax:CysVac2 i.t. delivery, and this was not observed after i.m. delivery of the vaccine. This is in agreement with other studies showing that IL-17 is strongly induced following mucosal vaccination and correlates with protection against *M. tuberculosis* infection[37, 38], yet the mechanism by which this protection is specifically mediated is not well defined. Blocking of IL-17 during *M. tuberculosis* challenge in CysVac2/Advax-vaccinated mice resulted in a complete loss of vaccine-induced protection and correlated with reduced recruitment of lung phagocytic cells such as neutrophils and macrophages, as well as a reduced priming of *M. tuberculosis*-specific T cells in the mLN (Fig 5, 6). Thus we can propose a mechanism (Fig 7) whereby IL-17 production by vaccine-specific CD4^+^ T_RM_ facilitates neutrophil and Monocytes/macrophages recruitment[11], and promotes the activation and proliferation of protective, pathogen-specific CD4^+^ T cells in the mLN^[39], [40],[41]^. Interstitial macrophages recruitment might be beneficial because these populations appear to possess significant antimycobacterial activity, as opposed to alveolar macrophages [42]. Previous work in a different infectious model has additionally demonstrated that IL-17 can mediates the recruitment of neutrophils with anti-bacterial potential[11]. Our observations differs to previous reports using the ID93+GLA-SE mucosal TB vaccine, which resulted in a Th17-dominated, tissue-resident response, but did not lead to improved protection when compared to parenteral immunization[43]. This may indicate that different adjuvants may induce diverse T_RM_ populations, and the capacity of Advax to direct a predominant, vaccine-specific T_RM_ response in the lung may be a key determinant in the protective efficacy of CysVac2/Advax vaccine.

**Figure 7.**
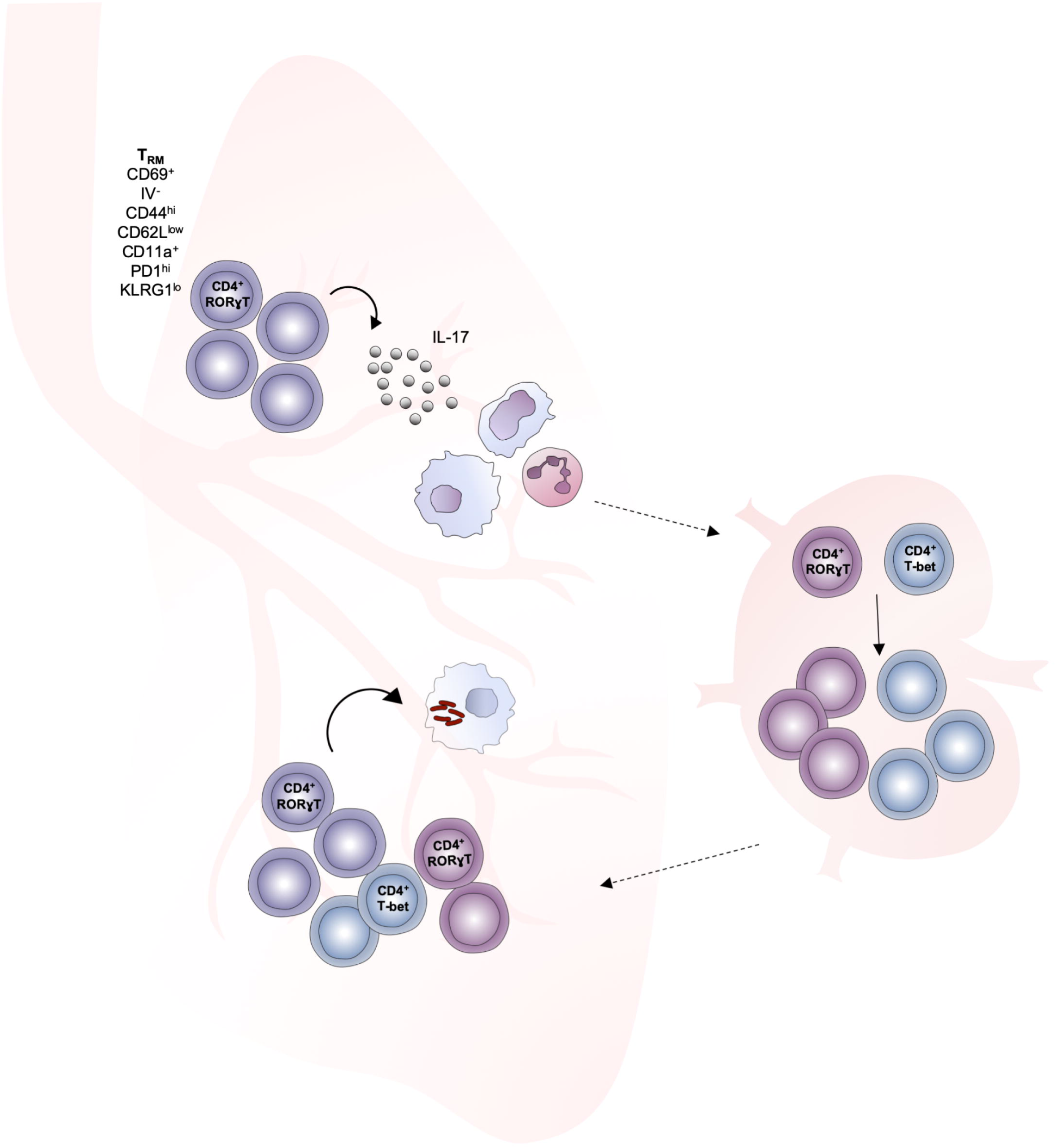
Proposed mechanism of immunity generated by pulmonary vaccination with CysVac2/Advax against *M. tuberculosis* infection. Pulmonary vaccination of CysVac2/Advax induces the development of a Th17 T_RM_ population of antigen-specific CD4^+^ T cells in the lung. Following aerosol *M. tuberculosis* infection, T cell IL-17 drives neutrophil and macrophage recruitment to the lung. This early response may be responsible for increased activation or trafficking of antigen presenting cells to the mLN, which in turn promotes the priming and proliferation of pathogen-specific CD4^+^ T cells in mLN that migrate to the lung and contribute to protection against *M. tuberculosis* infection.

Harnessing the protective role of IL-17 against pathogens needs to be balanced with potential tissue damage and pathology associated with excess cytokine levels[15]. Thus, adjuvants stimulating excessive levels of IL-17 are unlikely to be suitable for human lung administration, as observed for a candidate Sporotrichosis vaccine[44]. When CpG was removed from the CysVac2/Advax vaccine, the frequency of IL-17 secreting T cells in the lung was reduced, yet protection against *M. tuberculosis* challenge was not affected (Fig 2, S2 Fig). This suggests that a threshold of IL-17-secreting T cells may exist for vaccine-induced protection against *M. tuberculosis* and selecting the most ‘immunogenic’ vaccines based on the greatest level of effector responses may not be the best strategy for identifying an optimal *M. tuberculosis* vaccine formulation.

In conclusion, this report demonstrates that Advax-adjuvanted vaccines can be safely delivered to the lung to provide significant protection against aerosol *M. tuberculosis* infection. The protective effect of the vaccine was associated with targeted expansion of lung-resident T_RM_ and was dependent on IL-17 recruitment of phagocytic cells to the lung and enhanced priming of T cells in the mLN. As Advax-containing vaccines have proven safe and immunogenic in human trials against viral infection and allergy[19, 45, 46] and are safe and effective in pre-clinical trials as inhaled formulations[21], CysVac2/Advax is a promising candidate for assessment of safety, immunogenicity and efficacy as a pulmonary vaccine in human subjects.

## Materials and Methods

### Ethics statement

Female C57BL/6 (6-8 weeks of age) were purchased from Australian BioResources (NSW, Australia), and housed at the Centenary Institute animal facility (Sydney, Australia) in specific pathogen-free conditions. All mouse work was performed according to ethical guidelines as set out by the University of Sydney Animal Ethics Committee. All experiments within this manuscript were approved under protocol number 2017/011. University of Sydney Animal Ethics Committee guidelines adhere to the Australian Code for the Care and Use of Animals for Scientific Purposes (2013) as set out by the National Health and Medical Research Council of Australia.

### Bacterial strains

*M. tuberculosis* H37Rv (BEI Resources, USA) and BCG Pasteur were grown at 37° C in Middlebrook 7H9 medium (Becton Dickinson, BD) supplemented with 0.5 % glycerol, 0.02 % Tyloxapol, and 10 % albumin-dextrose-catalase (ADC) or on solid Middlebrook 7H11 medium (BD) supplemented with oleic acid–ADC.

### Mouse immunization, treatments and infection

CysVac2 fusion protein (Ag85B-CysD) was recombinantly expressed in *E. coli*, purified from inclusions bodies by ChinaPeptides (Shanghai, China), and refolded in Tris-buffer. Advax (delta-inulin, 50 mg/ml) and Advax^CpG^ (delta-inulin plus CpG, 50 mg/ml and 500 μg/ml, respectively) were provided by Vaxine Pty Ltd (Adelaide, Australia). Mice were anaesthetized by intraperitoneal (i.p.) injection of Ketamine/Xylazine (80/100mg/kg mouse) and then vaccinated with 1 mg of Advax or Advax^CpG^ and 3 μg of CysVac2 in a final volume of 50 μL PBS via i.m. route, using an insulin syringe (BD), or via the i.t. route, using PennCentury Microsprayer Aerosoliser (PennCentury, PA, USA). Three μg of CysVac2 alone or sterile PBS were administered as controls where appropriate. Mice were immunized subcutaneously with 5×10^5^ BCG for protection experiments.

For neutralization of IL-17, mice were injected i.p. with 250 μg of anti-IL-17A (clone TC11-18H10.1, Biolegend) or isotype control (clone RTK2071, Biolegend) one day prior to *M. tuberculosis* infection and then every three days for three weeks.

For *M. tuberculosis* challenge experiments, six or 8 weeks after the last vaccination mice were infected with *M. tuberculosis* H37Rv via the aerosol route using a Middlebrook airborne infection apparatus (Glas-Col, IN, USA) with an infective dose of approximately 100 viable bacilli. Three or 4 weeks later, the lungs and spleen were harvested, homogenized and plated after serial dilution on supplemented Middlebrook 7H11 agar plates. Colonies forming units (CFU) were determined 3 weeks later and expressed as Log_10_ CFU.

For intravascular staining of leucocytes, three minutes before euthanasia, mice were i.v. injected with 200 μL biotin-conjugated anti-CD45 mAb in PBS (15 μg/mL, Biolegend, clone 104) into the lateral tail vein. Detection of biotin was performed with APC-Cy7-conjugated streptavidin (BioLegend).

### Cell isolation, peptide stimulations and flow cytometry

PBMCs were isolated from whole blood as previously described[17]. Single cell suspensions were prepared from the lung as previously described[17]. PE-conjugated Ag85B_240-254_:I-A^b^ tetramer and APC-conjugated ESAT6_1-20_:I-A^b^ tetramer were provided by the NIH Tetramer Core Facility. For staining, cells were incubated with tetramers at 37 ^o^C for 1 hour. Cells were stained using the marker-specific fluorochrome-labeled mAbs indicated in S1 Table. To assess antigen-specific cytokine induction by T cells, PBMCs or single-cell suspensions from the lung were stimulated for 4 hours with CysVac2 (5 μg/mL) and then supplemented with brefeldin A (10 μg/mL) for further 10-12 hours. Cells were surface stained with Fixable Blue Dead Cell Stain (Life Technologies) and the marker-specific fluorochrome-labeled antibodies indicated in Supplementary Table. Cells were then fixed and permeabilized using the BD Cytofix/Cytoperm™ kit according to the manufacturer’s protocol. When required, intracellular staining was performed using mAbs against the specific cytokines (S1 Table). Samples were acquired on a BD LSR-Fortessa (BD), and analyzed using FlowJo™ analysis software (Treestar, USA). A Boolean combination of gates was used to calculate the frequency of single-, double- and triple-positive CD4^+^ T cell subsets. tSNE was run using default FlowJo parameters (iterations = 1000, perplexity = 30). Samples were randomly downsampled to 2000 events per sample and analysis was run on equal numbers of events per sample using FlowJo tSNE plugin.

### Statistical analysis

Statistical analysis was performed using GraphPad Prism 6 software (GraphPad Software, USA). The significance of differences between experimental groups was evaluated by one-way analysis of variance (ANOVA), with pairwise comparison of multi-grouped data sets achieved using the Tukey post-hoc test. Differences are considered statistically different when p ≤ 0.01.

## Acknowledgments

This work was supported by a National Health and Medical Research Council (NHMRC) Project Grant (APP1043519) and the NHMRC Centre of Research Excellence in Tuberculosis Control (APP1043225). We acknowledge the support of the European H2020 grant TBVAC2020 15 643381, the provision of reagents through the NIH Tetramer Core Facility and assistance from the Advanced Cytometry Facility and animal facility at the Centenary Institute. NP is supported by National Institutes of Health contract HHSN272201400053C and development of Advax adjuvant was supported by NIH Contracts AI061142 and HHSN272200800039C.

## Author contributions

### Conceptualization

Claudio Counoupas, Nikolai Petrovsky, Warwick J. Britton, James A. Triccas

### Data curation

Claudio Counoupas, Kia Ferrell, Anneliese Ashhurst, James A. Triccas

### Formal analysis

Claudio Counoupas, Kia Ferrell, Anneliese Ashhurst, James A. Triccas

### Funding acquisition

Nikolai Petrovsky, Warwick J. Britton, James A. Triccas

### Investigation

Claudio Counoupas, Kia Ferrell, Anneliese Ashhurst, Nayan Bhattacharyya, Gayathri Nagalingam

### Methodology

Claudio Counoupas, Kia Ferrell, Anneliese Ashhurst, Nayan Bhattacharyya, Carl G. Feng, James A. Triccas

### Project administration

James A. Triccas

### Resources

Carl G. Feng, Nikolai Petrovsky, Warwick J. Britton, James A. Triccas

### Supervision

Claudio Counoupas, James A. Triccas

### Writing – original draft

Claudio Counoupas, Kia Ferrell, James A. Triccas

### Writing – review & editing

Claudio Counoupas, Kia Ferrell, Anneliese Ashhurst, Nayan Bhattacharyya, Gayathri Nagalingam, Carl G. Feng, Nikolai Petrovsky, Warwick J. Britton, James A. Triccas

### Conflict of Interest Disclosure

NP is the research director for Vaxine P/L. All authors attest they meet the criteria for authorship.

---

**S1 Figure. Comparative analysis of multifunctional CD4^+^ T cell subsets before and after *M. tuberculosis* infection of mice vaccinated with CysVac2/Advax^CpG^ either via pulmonary or parenteral route.**

C57BL/6 mice (n=5-6) were vaccinated by either the i.m. or i.t. route with CysVac2 (CV2)/Advax^CpG^ (3 times, 2 weeks apart). One week after last vaccination mice were bled for vaccine immunogenicity assessment. Six weeks after last immunization mice were challenged with H37Rv by aerosol (~100 CFU) and four weeks later culled to enumerate bacterial burden and T cell phenotype in the lung. PBMCs (one week after last vaccination, panel a) and lung cells 4-weeks p.i. (panel b) were restimulated with CysVac2 fusion protein, and analysed for intracellular expression of IFN-γ, IL-2, IL-17, and TNF by flow cytometry. Boolean gating analysis was performed to identify subsets of CD4^+^ T cells expressing different combinations of these cytokines. Data is representative of 2 independent experiments. Significance difference between the groups was determined by ANOVA (*p<0.05; **p<0.01).

**S2 Figure. Comparative analysis of lung multifunctional CD4^+^ T cell subsets after *M. tuberculosis* infection of mice vaccinated via pulmonary route with either CysVac2/Advax or CysVac2/Advax^CpG^.**

C57BL/6 mice (n=5-6) were vaccinated by i.t. route with either CysVac2 (CV2)/Advax or CysVac2 (CV2)/Advax^CpG^ (3 times, 2 weeks apart). Six weeks after last immunization mice were challenged with H37Rv by aerosol (~100 CFU), and four weeks later culled to enumerate bacterial burden and T cell phenotype in the lung. Lung cells 4◻weeks p.i. were restimulated with CysVac2 fusion protein, and analysed for intracellular expression of IFN-γ, IL-2, IL-17, and TNF by flow cytometry. Boolean gating analysis was performed to identify subsets of CD4^+^ T cells expressing different combinations of these cytokines. Data is representative of 2 independent experiments. Significance difference between the groups was determined by ANOVA (*p◻<◻0.05; **p◻<◻0.01).

**S3 Figure. tSNE expression of markers in the lung of mice after *M. tuberculosis* infection and anti-IL-17 mAb treatment.**

Example of tSNE dimension 1 and 2 plots of the lung compartment show relative expression intensity of each indicated phenotypic marker. tSNE heat maps show fluorescent intensity of each marker for each event. Scales on the heat maps are individually generated for each surface marker from low to high expression. (Ashhurst, T. M. 2017, tSNEplots v1.3. GitHub repository).

**S4 Figure. Effects of blocking IL-17 during *M. tuberculosis* infection on CD4^+^ T cells proliferation in the lung.**

C57BL/6 mice (n=5-6) were vaccinated i.t. with CysVac2/Advax and treated i.p. with anti-IL-17 mAb, as described in Figure 5. Representative dot plot of the expression of Ki67 and RORγT on CD4^+^ T cells from the lung (a). Bar graphs showing numbers of total (b) and RorγT^+^ (c) proliferating CD4^+^ T cells enumerated in the lung. Representative dot plots show CD44 and either Ag85B:I-Ab or ESAT6:I-Ab staining on CD4^+^ T cells in the lung (d), with total number ± SEM of Ag85B:I-Ab^+^ (e) and ESAT6:I-Ab^+^ (f) CD4^+^ T cells in the lung. Data are pooled of 2 independent experiments. Significance of differences between the groups was determined by ANOVA (*p<0.05; ***p<0.001; NS=not significant).

**S1 Table. List of antibodies used.**

## Notes

#### Summary of Updates

Changed title and minor word changes in the text

