## Supplementary figures and images for "Mucosal delivery of a multistage subunit vaccine promotes development of lung-resident memory T cells and affords interleukin-17-dependant protection against pulmonary tuberculosis"

### S1

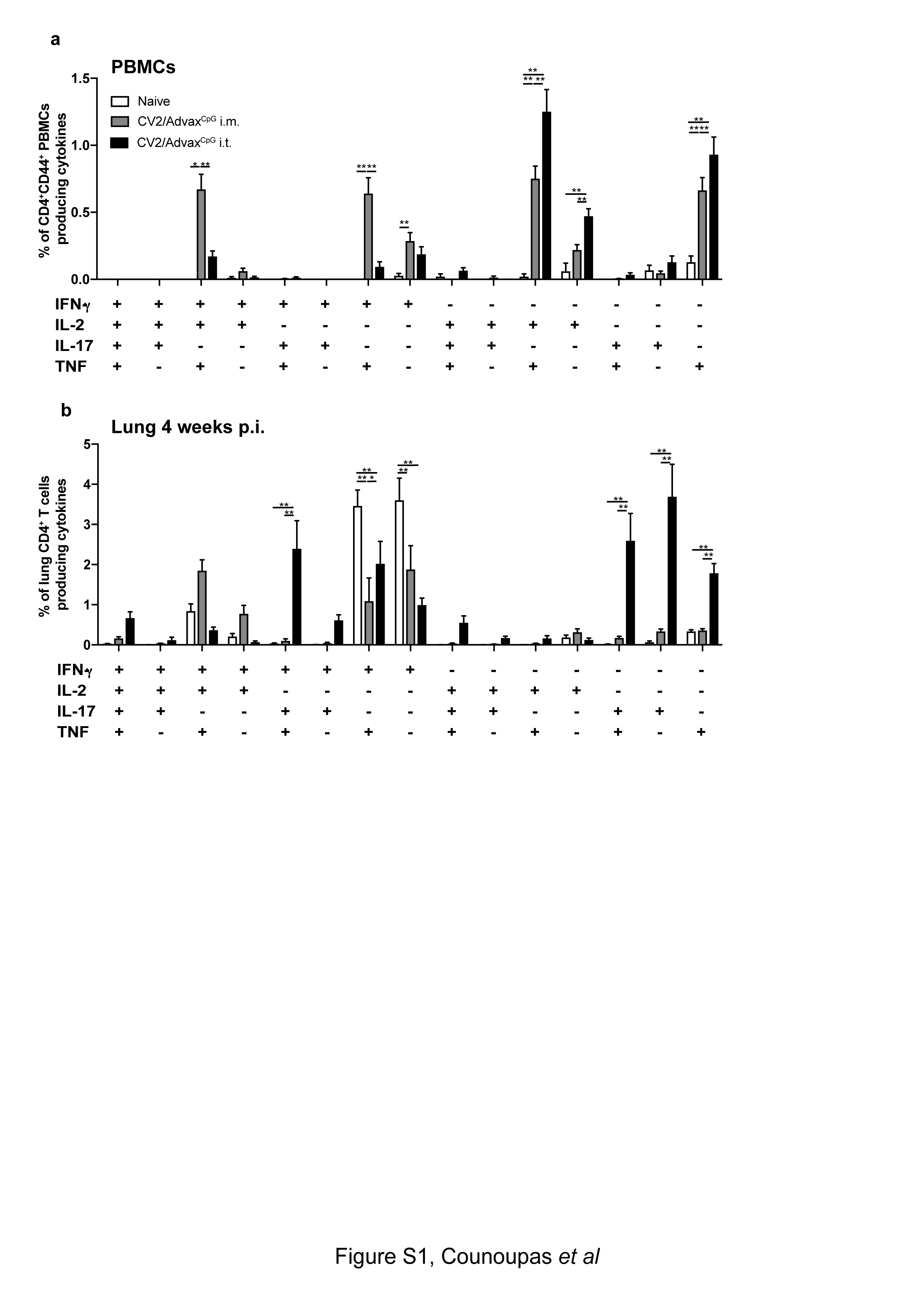

### S2

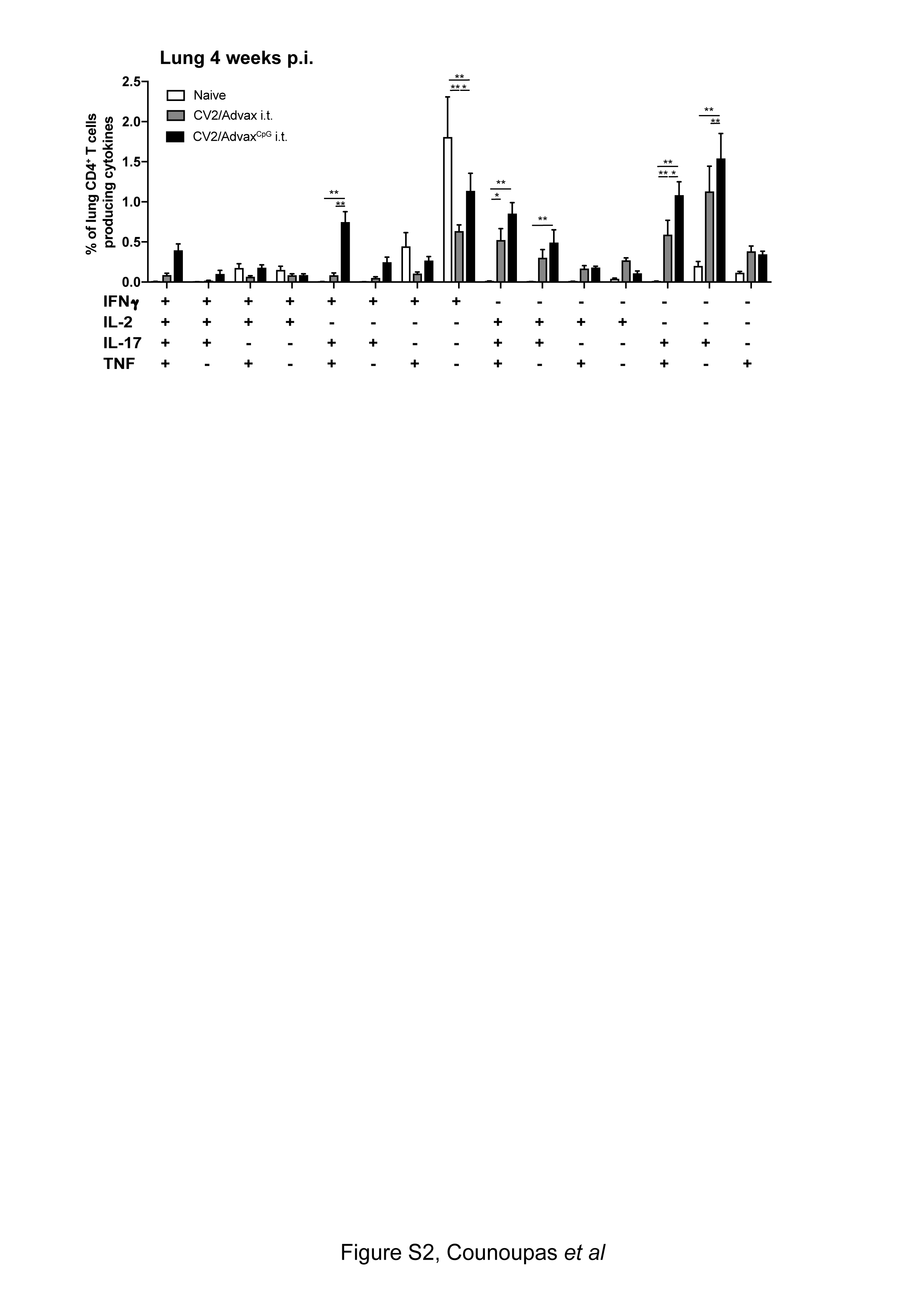

### S3

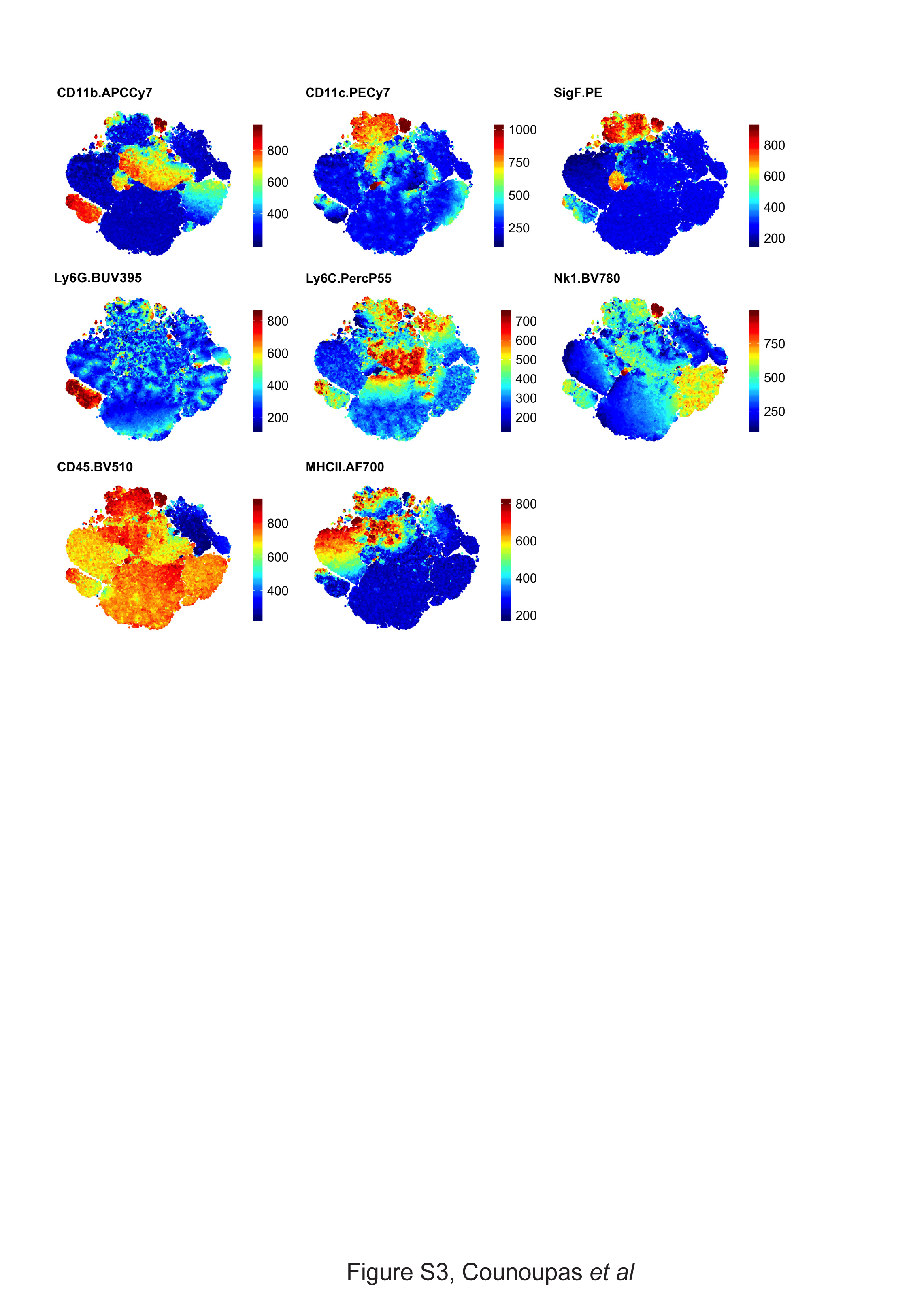

### S4

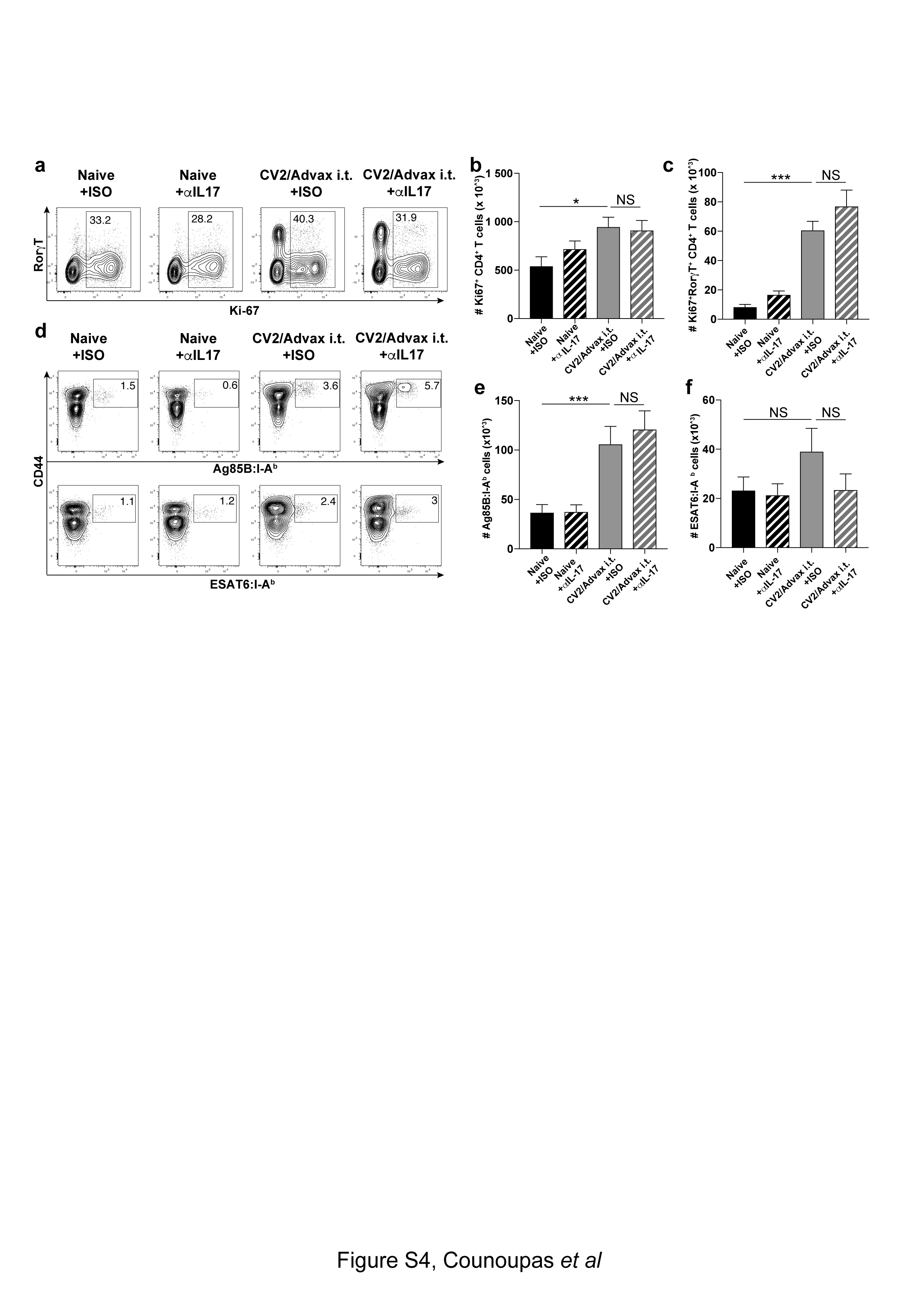
